# BaySiCle: A Bayesian Inference joint kNN method for imputation of single-cell RNA-sequencing data making use of local effect

**DOI:** 10.1101/2021.05.24.445309

**Authors:** Abhishek Narain Singh

## Abstract

There is a marked technical variability and a high amount of missing observations in the single-cell data that we obtain from experiments. Apart from that clearly each of the batch of experiments have a batch effect on every cell in the batch. This batch effect can be taken into advantage for dealing with imputation, given that all the cells in a given batch belong to the same tissue. Here we introduce ‘BaySiCle’, a novel Bayesian inference based method combined with k-nearest neighbors algorithm for the imputation of missing data in scRNA-seq counts. The priors are found out based on expression value across cells for all the single cells of the same batch. We demonstrate using sample scRNA-seq datasets and simulated expression data that BaySiCle allows robust imputation of missing values generating realistic transcript distributions that match single molecule fluorescence in situ hybridization measurements. By using priors as obtained by the dataset structures in the not just the experimental set-up batch, but also the same group of cells, BaySiCle improves accuracy of imputation to be that much closer to its similar alternatives.

**Availability and implementation:** The Python Jupyter notebook ‘BaySiCle’ is published on GitHub GitHub - abinarain/BaySiCel: Single Cell Data Imputation using Bayesian statistics and kNN

## Introduction

Single-cell RNA-sequencing, scRNA-seq, has recently emerged as a novel method of choice for profiling gene expression heterogeneity across tissues in health & disease (Baslan and Hicks, 2017; Chen et al., 2018) and also for other metabolic profiling. However, as the technique relies on the detection of minute amounts of RNA content of one single cell, scRNA-seq is many times the value which is read and is highly prone to technical biases. The technical reason for this is that scRNA-seq library preparation protocols recover only a small fraction of the total RNA molecules present in every single cell. The ‘dropouts’ or the zero values are generated for many genes and what we get is a sparse matrix. Another term, ‘capture efficiency’, is used to describe the amount of genes for which the expression level values are obtained. Besides the above mentioned points, the expression values have a confounding effect known as batch effect which according to some researchers are a major problem (Bacher and Kendziorski, 2016; Vallejos et al., 2017; Ziegenhain et al., 2018). The origin of batch effects is not completely understood but the differences in average capture efficiencies across experiments has an effect on it (Hicks et al., 2018). Many of the recent studies have suggested that the data be normalized first (Wenhao Tang et. al., 2020) to take care of batch effect before going for further processing. However, in this paper, we propose that the batch effect can be taken to an advantage for imputation such as by Bayesian inference followed by kNN, and any normalization that should be done, can be done after the imputation has been done using the techniques and/or codes as in this paper in the form of ‘BaySiCle’. The method of imputation also acts as a regularizer for a model as has been demonstrated by Bayesian clustering approach as used in Melissa (Kapourani CA et. al.). Among other recent methods applied for imputation has been generative adversarial networks such as for the tool scIGANs (Xu et. al.). netImpute employs Random Walk with Restart (RWR) to adjust the gene expression level in a given cell by borrowing information from its neighbors in a gene co-expression network (Zand et. al.). Iterative imputation approach based on efficiently computing highly similar cells method has been used (Moussa et. al.) and in line with this BaySiCle uses a similar concept by imputing the cells in the same batch. Badsha et. al. made use of an autoencoder neural network for single-cell gene expression. Another method is based on K-nearest neighbor (kNN) smoothing, and uses Poisson distribution and aggregate information from similar cells (Wagner et al., 2018). In BaySiCle, I have made use of SCEDAR Python library (Zhang et. al.) for using the k-nearest neighbor for imputation. The (kNN) method essentially takes care of the locality effect, much of an alternative to Moussa et. al. Although we have used SCEDAR for demonstration purposes of how kNN can be used, future implementation of BaySiCle can well use other standard libraries for the purpose such as the Scikit-learn.

## Materials & Methods

Sample RNA-Sequencing data was provided with courtesy by Dr Ville Hautamäki who is the author of paper (Trung Ngo Trong et al., 2020), in which Bayesian inference method has also been used to get the latent values. However, in that paper, since the cells coming from a common group are not taken into account, they make use of another distribution (Kingma and Welling, 2014) to get the posterior probability. Our method of Bayesian inference to get the posterior probability for the values of gene expression which needs imputation relies a lot on the batch or group effect, and so the formula is designed accordingly unlike in Trung Ngo Trong et al., 2020, which did not take batch effect into account for their advantage. The sample data is also provided in the GitHub link specified to download the Python script.

The count of each gene in each cell follows a Poisson–gamma mixture, also known as a negative binomial model can be used. However, given that we do not know much of relations between the genes, and we do know much of relations between the cells, it would make sense to use cells as evidence in the Bayes theorem. So the Posterior probability would be:

### p(GENEi | CELLj) = p(CELLj|GENEi) p(GENEi) / p(CELLj)

Here p(CELLj|GENEi) is likelihood, p(GENEi) is prior probability, and p(CELLj) is evidence, as per the terminology used in Bayesian inference.

This posterior probability p(GENEi|CELLj) needs to be multiplied by the MEAN of the expression level of GENEi in order to convert the probability value which is between 0 and 1, to a gene expression value.

### Final Score to be imputed = Posterior Probability (X) MEAN Expression Score of GENEi

The prior probability p(GENEi) is known for any gene = 1/ total number of genes examined = 1 / No. of columns in the data sheet where each column is for a unique gene expression value

### The EVIDENCE of p(CELLj) = (Total number of cells similar to CELLj) / (Total number of similar cells groups)

For extracting the total number of similar cells groups, the naming convention of the cell was followed such that programmatically we can put a condition in loop which parses the data file, such that the first word of cell is the same depicting same batch and the first letter of second word being same implying same or similar cell i.e. of same group. We also leave the responsibility of sorting the data as per the batch and group order such as alphabetically to the user before processing the data by this script. Note how cleverly we have differentiated cells to be of not just belonging to the same batch but also to the same group to ensure that the cells examined for imputation are as similar as possible. As an example cells with Id VZA00602.A03, VZA00602.A05 are of the same batch VZA00602 as well as from the same group ‘A’. Cells with Id VZA00602.B03 and VZA00602.A03 are from the same batch but not the same group. Cells with Id VZA00612.A03 and VZA00602.A03 are totally different given that they are from different batches.

Thus, once we have the cells and their counts calculated, in order to get likelihood value in the Bayesian inference formula P(CELLj|GENEi), we have to look for the counts entries where the GENEi value is non-zero. The Python script works by first conducting a Bayesian Posterior probability calculation wherever possible to impute the missing values. Thereafter, if there are still any missing values still remaining (also called ‘dropouts’) as shown by 0s are imputed using kNN algorithm, as discussed earlier. It should be noted that those genes which did not yield any value in any of the single-cell data were dropped out completely and not imputed at all. Table 1 shows a snippet of sample data for scRNA-seq that was imputed. Notice how sparse the matrix is, as this is how typically the scRNA-seq data looks like.

**Table 1:** Snippet of Data showing the cell name and the genes with their expression values

|  | Cell | A1BG | A1BG-AS1 | A2M | A2M-AS1 | A2MP1 | A3GALT2 | A4GALT | AAAS | AACS | AAED1 | AAGAB | AAK1 | AAMDC | AAMP |
| --- | --- | --- | --- | --- | --- | --- | --- | --- | --- | --- | --- | --- | --- | --- | --- |
| 0 | VZA00602.A03 | 0 | 0 | 1289 | 1281 | 0 | 0 | 0 | 0 | 0 | 0 | 152 | 0 | 0 | 684 |
| 1 | VZA00602.A05 | 0 | 0 | 1051 | 1450 | 0 | 0 | 0 | 0 | 0 | 0 | 0 | 0 | 0 | 681 |
| 2 | VZA00602.A07 | 0 | 0 | 1640 | 1805 | 0 | 0 | 0 | 0 | 0 | 0 | 0 | 0 | 0 | 1074 |
| 3 | VZA00602.A11 | 0 | 0 | 1400 | 1090 | 0 | 0 | 0 | 0 | 0 | 0 | 0 | 0 | 0 | 661 |
| 4 | VZA00602.A15 | 0 | 0 | 1216 | 2479 | 0 | 0 | 0 | 0 | 0 | 0 | 549 | 0 | 0 | 764 |

## Results

t-SNE (Roweis et al., 2002) 2-D plots are generated before and after the imputation that give a comparison of the impact of imputation. First we see the plot without any imputation as in figure A below. The figure is plotted without any label at the moment.

**Figure A:**
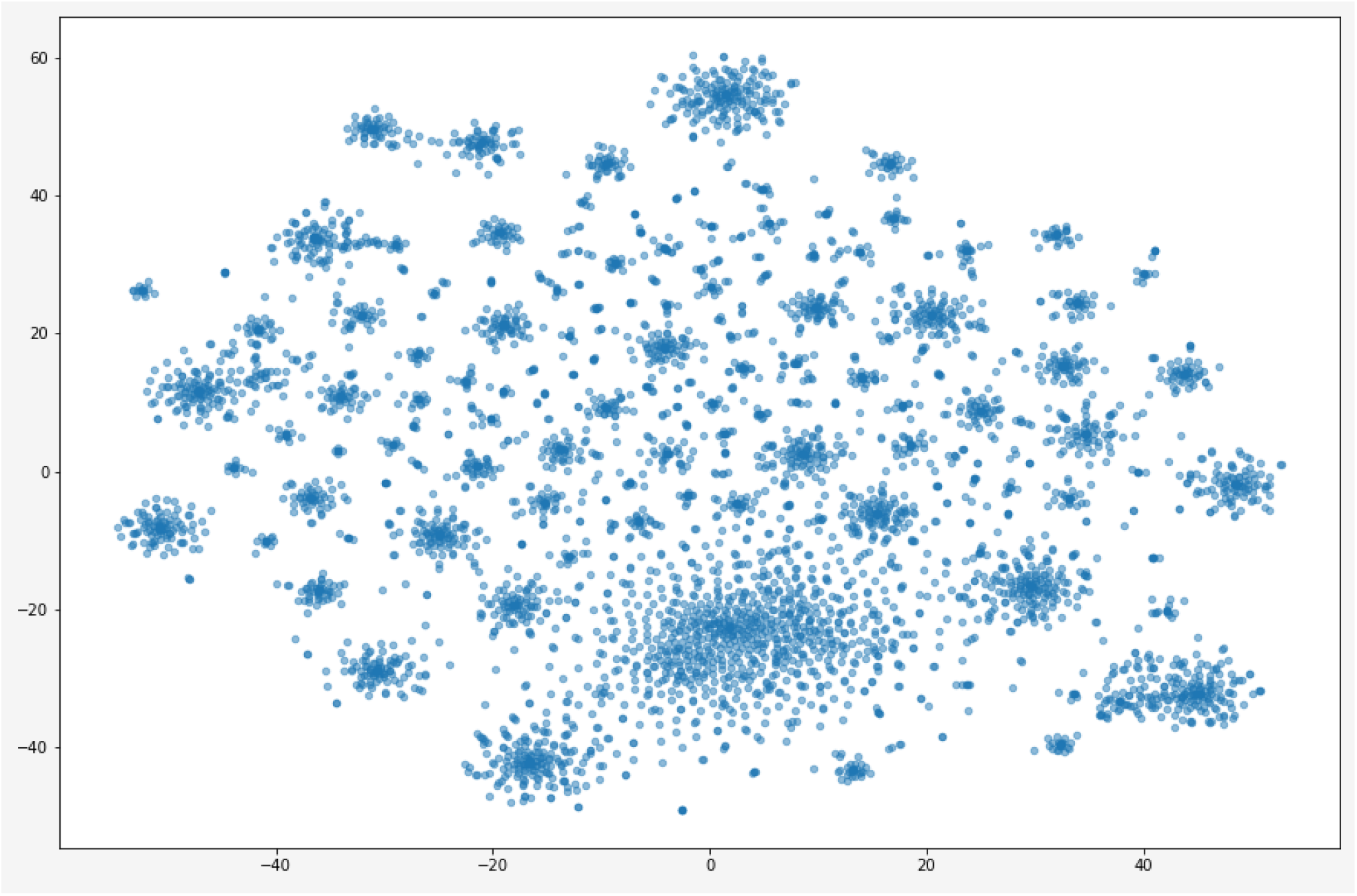
tSNE 2D plot before imputation.

A possible label for the plots could have been the group name or the batch name, which we leave for future exploration. At the moment, we would like to see the plots for qualitative purposes. After applying the Bayes inference using the group method as discussed above, we see a reduction of ‘dropouts’ from 1446458 to 910073. Difference 536,385 ZEROS have been imputed.

### Percentage of cells imputed using BAYES theorem = 536385 / 1446458 *100% = 37%

The remaining ‘dropouts’ were imputed using the kNN method with k=30. Eventually, we can then see our new tSNE plots as in figure B, and we note it to be significantly different than that of Figure A which was not imputed.

**Figure B:**
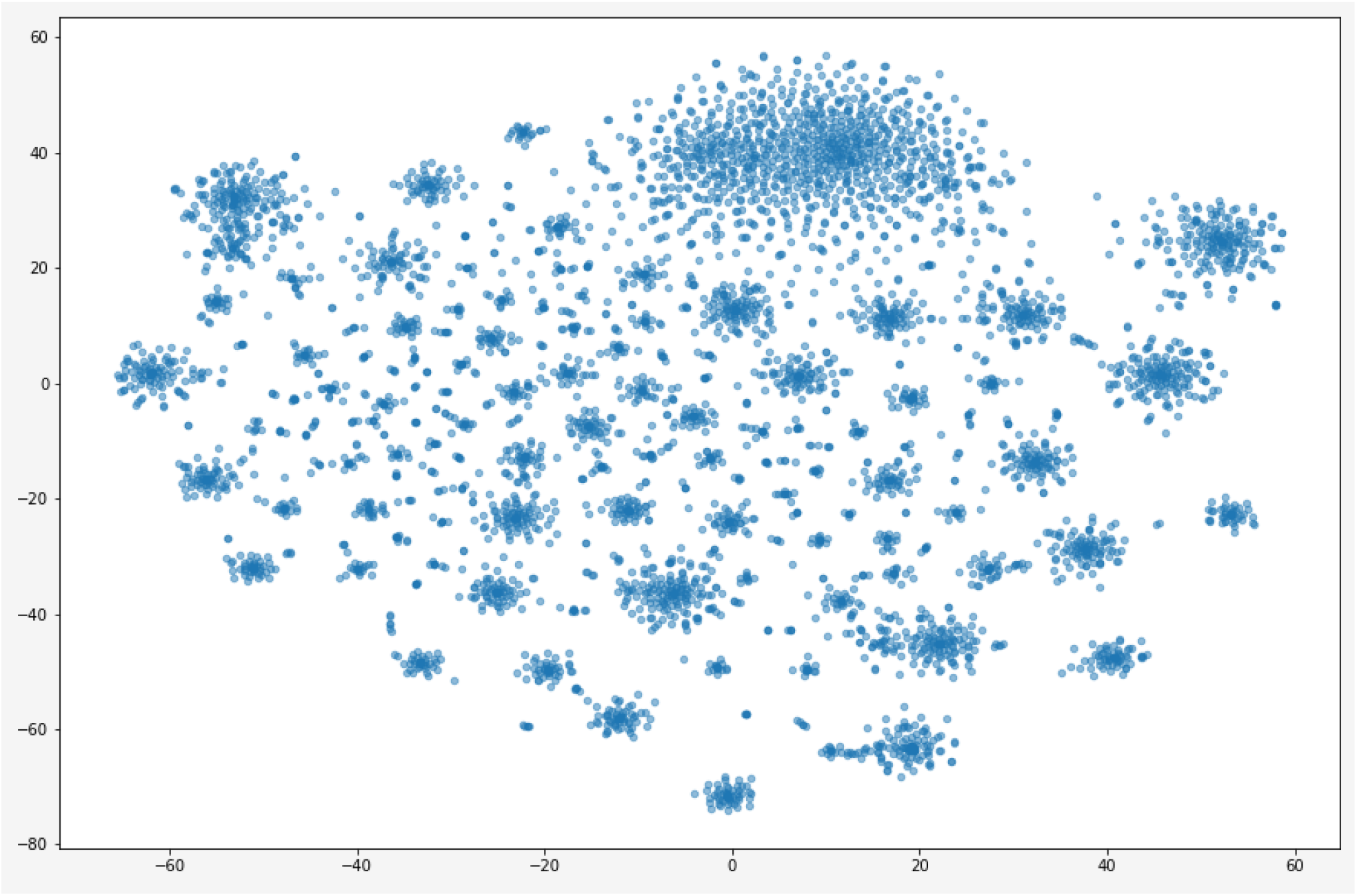
tSNE plot of scRNA-seq data after Bayesian group inference and kNN imputation of the dropouts.

In order to further qualitatively realize that this is different from a simple imputation by kNN method, we also did a full imputation of all the dropouts using kNN keeping k-30, and note that the plot in Figure C is significantly different than that of Figure B.

**Figure C:**
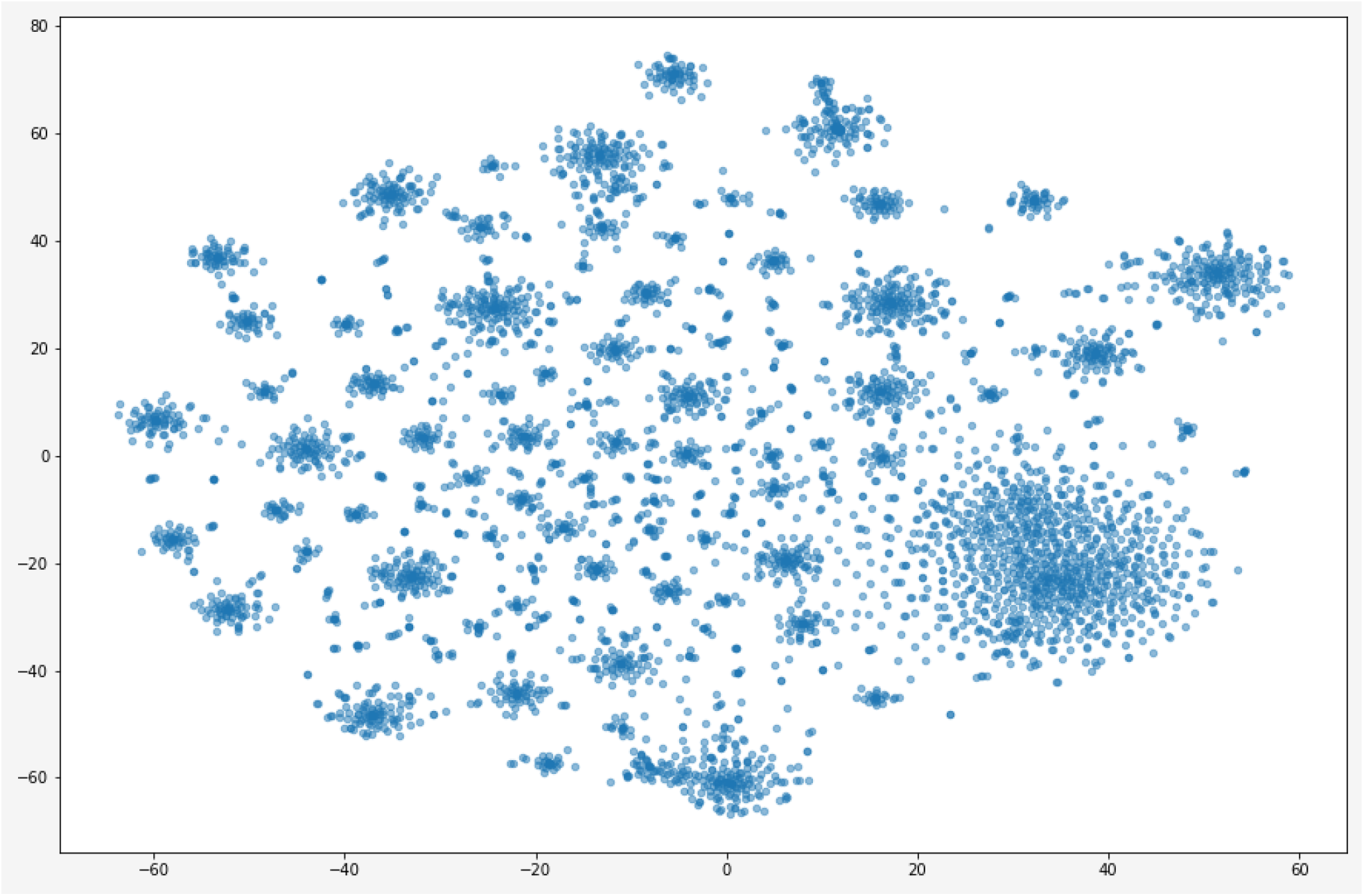
tSNE 2D plot of kNN imputed scRNA-seq data Conclusion & Future Work

In this article, we introduced a method of bayesian inference taking advantage of locality of the dropouts based on Bayesian posterior probability using the knowledge of the group of cells instead of bulk data. The locality aspect of the remaining of the un-imputed data using Bayesian approach is then taken care of by kNN method. We believe that the combination of these two locality based imputation methods and in the described order can well relate the true value of the scRNA-seq data. What can be done in the future is that the imputed values can then be normalized. Also, in future, it would make sense if we carry out a comparative performance analysis in terms of the results obtained from BaySiCle compared to the other tools that are out there. Clearly, plotting the t-SNE 2D plots with an appropriate label could be also more informative which is planned for in the future. A possible incorporation of deep learning techniques can also be explored in the future for improvising BaySiCle. Clearly, making the Python Jupyter notebook code as on GitHub link converted to automated software tool also remains one of the tasks in future agenda, although for all practical purposes, following the steps on the notebook would typically lead to the imputation as described in this paper.

## Notes

https://github.com/abinarain/BaySiCel

### Competing Interest Statement

The authors have declared no competing interest.

